# Metabolic Signatures of Immune Checkpoint Inhibitor Response in Gynecologic Cancers: Insights from Flux Balance Analysis

**DOI:** 10.1101/2025.01.14.631240

**Authors:** Gideon Idumah, Lin Li, Lamis Yehia, Haider Mahdi, Ying Ni

## Abstract

Modifiers of immune checkpoint inhibitor (ICI) responses in cancer patients are complex and remain poorly characterized, especially in gynecologic cancers. In this study, we explored fluxomic biomarkers that differentiate responders from non-responders to ICIs in a series of 49 patients with gynecologic cancers, including ovarian, cervical, and endometrial cancers. By applying metabolic enzyme expression as constraints, we utilized an objective-customizable flux balance analysis within a genome-scale metabolic model to predict the metabolic flux differences between responders versus non-responders of ICI treatment. We identified three reactions with consistent differential activity across all ten different optimization objectives: Succinate Dehydrogenase (SUCD1m) in the citric acid cycle, NADH: Guanosine-5-Phosphate Oxidoreductase (r0276) involved in purine catabolism, and Ornithine Transaminase Reversible, Mitochondrial (ORNTArm) in the urea cycle. Additionally, reactions within the folate cycle subsystem, particularly involving MTHFD2, demonstrated significance in distinguishing treatment responses, aligning with previous findings linking MTHFD2 to immune evasion and tumor progression. To further analyze the association between metabolic features and survival outcomes, we implemented machine learning models that integrate multi-omics data. Our model included clinical-pathologic, molecular-genomic features (gene expression, TGF-β score, immune cell abundance from transcriptomic deconvolution), and significant reaction fluxes. Our findings suggest that SUCD1m, MTHFDm and ORNTArm are important metabolic biomarkers that could serve as predictive indicators for ICI response and, if validated in a larger cohort, may guide the development of targeted therapies to enhance treatment efficacy for gynecologic cancer patients. This study highlights the use of genome-scale metabolic modeling to identify clinically relevant biomarkers and improve therapeutic strategies.

## 1. Introduction

Immunotherapy with immune checkpoint inhibitors (ICI) has emerged as a promising option in solid tumors like lung and urothelial cancers, as well as melanoma^1^. However, this approach has yielded limited success in treating gynecologic cancers, with response rates in recurrent scenarios hovering between 11% and 17%^2–4^. A crucial field of research now focuses on figuring out how to improve treatment efficacy and better understand mechanisms driving resistance to ICI in gynecologic cancers and identify the subset of patients who will benefit from immunotherapy.

The metabolic environment of the tumor microenvironment (TME) plays a critical role in determining treatment outcomes. It is well established that tumor cells reprogram their metabolism to support rapid growth and proliferation^5^, leading to increased competition for nutrients between tumor and immune cells and the creation of an environment that inhibits immune activity. This dysregulation of metabolic pathways within immune cells, as well as changes in the number or composition of nutrients and metabolites in the TME, are now emerging as important factors that regulate the body’s immune response against tumors^6–8^. This in turn affects the efficacy of immune-based treatments, including ICI, by reducing T cell infiltration and activity within the TME.

However, knowledge about the utilization of metabolic pathways requires quantification of metabolic fluxes (i.e., the rate at which a substance is transformed into another through a given reaction or pathway)^9^. As quantification of the full complement of cell metabolites (metabolomics) alone does not provide information on internal fluxes^10^, a more functional approach using computational models of metabolic networks is necessary. This approach offers a unique way of analyzing entire metabolic networks at a systems level, considering all interconnected biochemical reactions simultaneously, which may uncover novel metabolic pathways, regulatory mechanisms, and metabolic interactions that may not be apparent from metabolomics data alone. A well-accepted computational approach for analyzing metabolic networks is by using Genome-Scale Metabolic Models^11^ (GSMM). GSMMs are comprehensive computational models that represent the entire set of metabolic reactions occurring within an organism, typically at the genome level. These reconstructions integrate information about the genome sequence, enzymes, associated gene-protein-reaction (GPR) rules, metabolites, biochemical pathways, and known metabolic reactions to provide a systematic and quantitative framework for studying cellular metabolism.

To date, no study has developed personalized genome-scale models to investigate metabolic differences and system-wide alterations in redox metabolism among GYN cancer patients received immunotherapy. In this study, we address this gap by developing a unique computational methodology to predict reaction fluxes and to investigate differences in co-factor production between responders and non-responders to ICI in a series of patients with gynecologic cancers. We explored the relationship between the predicted metabolic fluxes and some important immunosuppressive biomarkers identified in our previous studies^1,12^, such as TGF-β and *CD47* which were reported to be associated with lower response to ICI. Alongside with machine learning models, we associated these fluxes and multi-omics biomarkers with resistance to ICI. This work offers insights into the metabolic characteristics linked to ICI response and suggests possible multi-omics indicators to enhance patient selection for immunotherapy.

### 2. Materials and methods

## 2.1. Study Cohort

We collected clinical and pathological data from 49 patients with ovarian, endometrial, and cervical cancers at various stages, all treated with the immune checkpoint inhibitors, nivolumab, pembrolizumab or avelumab. The clinic-pathological summaries of these patients can be found in supplementary table 1. Patients were categorized as responders if they demonstrated a reduction in radiologic tumor burden, meeting criteria for partial or complete response based on RECIST 1.1, as assessed by CT scans per judgment of their treating physician. To qualify, patients were required to complete at least two cycles of immunotherapy with interval imaging to assess response compared to pre-treatment imaging. Information on the type and frequency of immune-related toxicities were retrospectively gathered from the patient’s medical records. The progression-free survival was calculated from the time of start of immunotherapy with immune checkpoint inhibition to disease progression, last follow up or death. Overall survival was calculated from the time of start of immunotherapy with immune checkpoint inhibition to death or last follow up. This was performed to elucidate the impact of immunotherapy on these survival endpoints.

## 2.2. RNAseq data

Formalin fixed paraffin embedded (FFPE) specimens with sufficient tumor content (>20%) were sequenced by MedGenome (Foster City, CA). All 49 samples passed RNA extraction and library prep quality checks and underwent mRNA sequencing. Low-quality sequence reads were excluded, and high-quality reads were trimmed and processed using FastQC (v0.11.8). Paired-end reads were aligned to the reference human genome (GRCh37/hg19) using STAR (v2.7.3a), and gene expression was estimated with HTSeq (v0.11.2), counting only reads uniquely mapped to single genes. More information can be found in our previous publication^1^.

## 2.3. TGF-β score and immune cell abundance estimation

A detailed description of how the TGF-β score was calculated can be found in our previous publication^1^. Briefly, the TGF-β score was generated by applying single-sample enrichment analysis to 6 genes (*SLC20A1, XIAP, TGFBR1, BMPR2, FKBP1A, and SKIL)* regulated by the TGF-β signaling pathway that had lower expression in patients who responded to immunotherapy. On the other hand, immune cell abundance estimation was based on LM22^13^, which is a signature matrix file consisting of 547 genes that accurately distinguish 22 mature human hematopoietic populations isolated from peripheral blood or in vitro culture conditions, including seven T-cell types, naïve and memory B cells, plasma cells, NK cells, and myeloid subsets. Expression levels for each LM22 cell type were calculated using the rank-based single-sample gene set scoring method as described by Foroutan et al^14^.

## 2.4. Genome Scale Metabolic Model

Genome-scale network reconstructions are built from curated and systematized knowledge that enables them to quantitatively describe genotype-phenotype relationships^15,16^. In the case of metabolism, by mapping the annotated genome sequence to a metabolic knowledge base such as KEGG, one can reconstruct a metabolic network composed of all known metabolic reactions. Such genome-scale reconstructed networks represent organized and systematized knowledgebases that provide a comprehensive understanding of cellular metabolism across various microorganisms and some mammalian tissues, incorporating all known metabolic genes and reactions based on genome annotations and experimental evidence (represented example as in Figure 1A). These reconstructed networks have multiple uses, including conversion into computational models that interpret and predict phenotypic states and the consequences of environmental and genetic perturbations.

**Figure 1:**
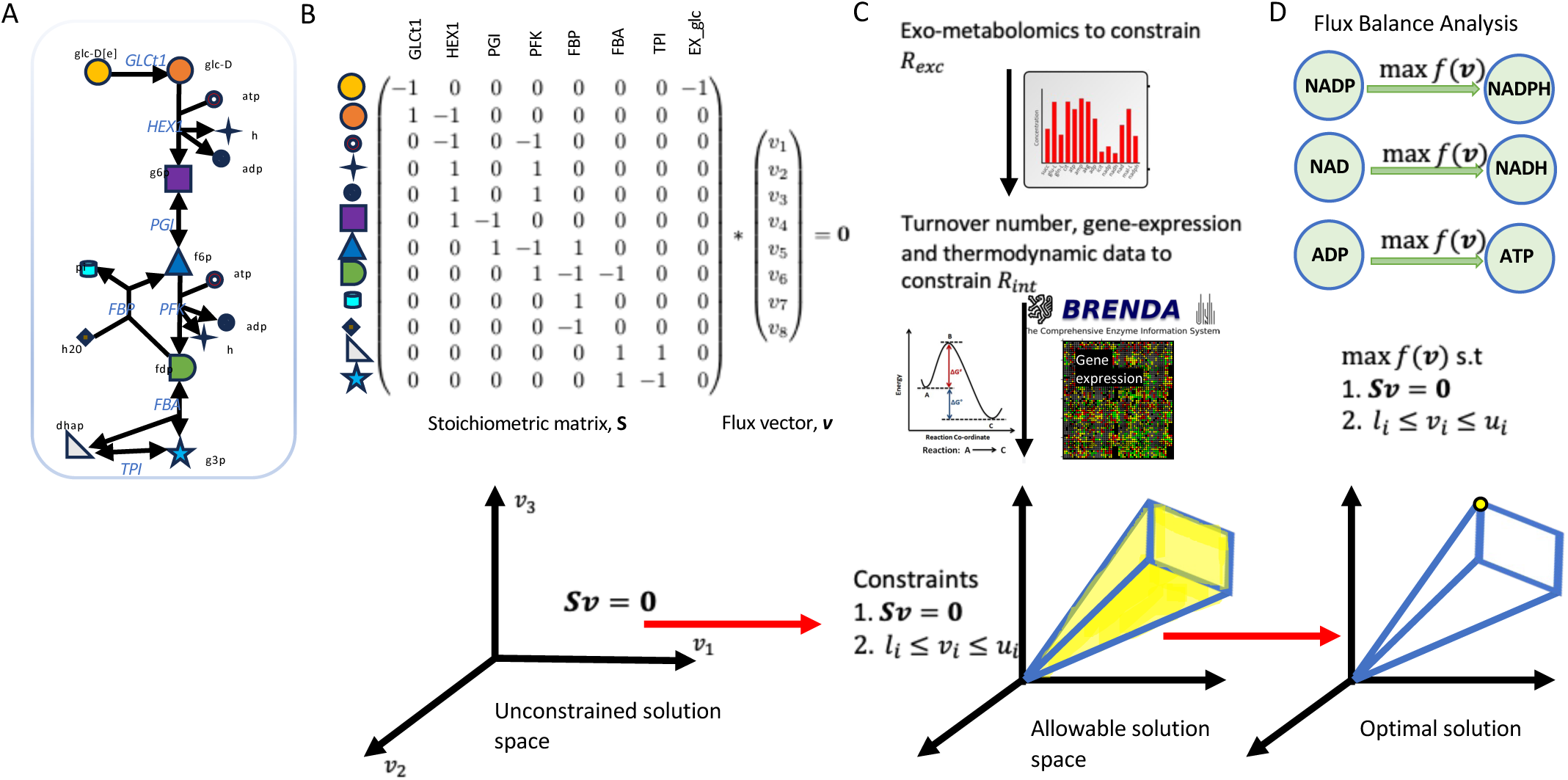
Conceptual framework of constraints-based modeling. **(a)** Depiction of the first few reactions of glycolysis to illustrate the concept of GSMM. **(b)** Stoichiometric matrix (S) corresponding to these reactions. At steady state, Sv = 0, where *v* represents the flux vector. Without constraints, flux distribution may lie at any point in the solution space. **(c)** Incorporating constraints such as turnover numbers, gene expression, and thermodynamic data, refines the solution space to allowable flux distributions. This limits the possible flux distributions to within the constrained space, excluding any points outside the defined bounds. **(d)** Optimizing an objective function identifies a single optimal flux distribution at the edge of the allowable solution space.

The metabolic network in a genome-scale metabolic reconstruction can be converted into a mathematical format — a stoichiometric matrix (**S**) of size *m x n* — where *m* the number of metabolites and *n* represents the number of reactions. As seen in Figure 1B, each entry is the corresponding coefficient of a particular metabolite participating in a reaction such that S_ij_ < 0 for every metabolite consumed and S_ij_ > 0 for every metabolite produced. A coefficient of zero is used for every metabolite that does not participate in a particular reaction. Therefore, **S** is a sparse matrix because most biochemical reactions involve only a few different metabolites. The flux through all of the reactions in a network is represented by the vector **v**, which has a length of *n*.

The resulting genome-scale model is a mathematical representation of the reconstructed network that facilitates computation and prediction of multi-scale phenotypes through the optimization of an objective function of interest. These models have been developed for a wide variety of organisms, including humans. For our work, we utilize the community curated Recon3D^17^ as our generic human genome-scale metabolic model. Recon3D represents the most comprehensive and up-to-date human metabolic network model, accounting for 3,288 genes, 13,543 metabolic reactions involving 4,140 unique metabolites, and 12,890 protein structures.

### 2.4.1. Flux balance analysis

There are various methods for simulating metabolism in a GSMM. Examples include Flux Balance Analysis (FBA), Flux Variability Analysis (FVA), Minimization of Metabolic Adjustments (MOMA), dynamic FBA (dFBA), and others. Flux balance analysis (FBA) is preferred for this work because it can predict steady-state flux values through an entire metabolic network with thousands of reactions within seconds^18^. By combining the stoichiometric representation of the human metabolic network, constraints on the fluxes through metabolic reactions, and an objective function to maximize a particular metabolic phenotype, predictions of maximum reaction fluxes or metabolite production rates under physiological constraints are generated (Figure 1C-D). In contrast to the traditionally followed approach of metabolic modeling using coupled ordinary differential equations, FBA requires very little information in terms of the enzyme kinetic parameters and concentrations of metabolites in the system. It achieves this by making two assumptions, steady state and optimality. The predictive power and computationally inexpensive nature of FBA have led to its use in a variety of different biomedical areas, including drug target identification^19,20^, radiation-resistance^21^, and human disease modeling^22–25^.

The first assumption is that the modeled system has entered a steady state, where the metabolite concentrations no longer change, i.e., in each metabolite node the producing and consuming fluxes cancel each other out. This implies that **Sv = 0** The second assumption is that the organism has been optimized through evolution for some biological goal, such as optimal growth or conservation of resources.

FBA seeks to maximize or minimize an objective function **Z = c^T^v**, which can be any linear combination of fluxes, where **c** is a vector of weights indicating how each reaction contributes to the objective function. In practice, when only one reaction is desired for maximization or minimization, **c** is a vector of zeros with a value of 1 at the position of interest. A key part of FBA is the ability to add constraints to the flux rates of reactions within networks, forcing them to stay within a range of selected values. This enables the model to more accurately simulate real metabolism. In summary, flux balance analysis can be represented as a Linear Programming (LP) optimization problem:

Maximize: **c^T^v** subject to:

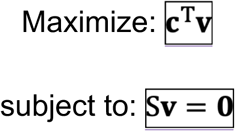

and

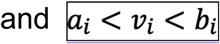

The output of FBA is a particular flux distribution **v,** which maximizes or minimizes the objective function. We used the CPLEX^26^ optimization software to solve these FBA LP problems.

### 2.4.2. Objective functions

While maximizing biomass production is often used as the primary objective function in GSMMs, cells could have different or multiple objectives depending on environmental conditions, growth phases and metabolic needs. Hence, it might be useful to explore other forms of objective functions that cells might prioritize in addition to biomass production.

To maximize the production of a particular metabolite in a metabolic network, an artificial demand reaction can be added to the network, and the flux through this new objective function can be maximized. This in turn maximizes the fluxes through other reactions through the metabolic network that produces the metabolite. For example, to maximize the production of NADH in the mitochondria, the objective function would be:

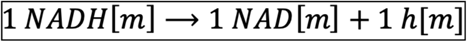

Given our interest in redox metabolism, we formulated several objective functions to assess differences between responders and non-responders to immune checkpoint inhibitors among our cohort. In addition to biomass production, these functions aimed to maximize the production of redox co-factors such as NADH and NADPH in the cytoplasm, mitochondria, and across all compartments where they are involved. We also considered the objective of maximizing ATP production. The ten objective functions implemented were as follows: biomass production; NADP (cytoplasmic), NADP (mitochondrial), NADP (all compartments), NADPH (cytoplasmic), NADPH (mitochondrial), NADPH (all compartments), ATP (cytoplasmic), ATP (mitochondrial), ATP (all compartments).

## 2.5. Context-specific model

A significant challenge in genome-scale modeling is the difficulty in assigning constraints to reaction fluxes. Metabolic networks are often distributed as reconstructions and not as condition-specific models. As a result, exchange and internal reactions are typically unconstrained, i.e., they have “infinite bounds”. Moreover, these reconstructions represent a superset of potentially inactive reactions, as cells selectively utilize reactions based on specific needs. To address this, it is important to incorporate as much relevant information as possible to constrain the reaction fluxes. This includes integrating transcriptomics, genomics (if available), kinetics, and thermodynamic data to establish quantitative constraints on the maximum flux and directionality of metabolic reactions.

To develop a context-specific model, we adapted the methodology outlined by Lewis et al^21^. Their approach incorporated transcriptomic and mutational data, along with genome-scale kinetic and thermodynamic parameters, to define quantitative constraints on metabolic fluxes in the generic human Recon3D reconstruction. We followed their procedure to address missing and inaccurate redox-based reaction information within Recon3D, with a few modifications specific to our study:

1. We removed the neomorphic 2-hydroxyglutarate-producing reaction catalyzed by mutant IDH, as well as the associated export reactions, because we do not have *IDH1* mutation data and are not considering this mutation in our work.
2. We added back oxalosuccinic acid-producing IDH1 reactions – specifically “r0423”, “r0424”, “r0425” and “r0422” – which were blocked in their model.
3. We identified and removed duplicate nadph[n] and nadp[n] metabolites.

Subsequently, we updated the Stoichiometric matrix, metabolite list and corresponding reaction formulas.

To set upper flux bounds for reactions in Recon3D that have associated Gene-Protein-reaction (GPR) rules and Enzyme Commission (EC) numbers, we used the formula:

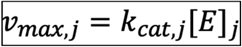

where *k_cat,j_* is the turnover number (in units of *hr*^−1^and [*E*]*_j_* is the enzyme abundance (in units of mmol gD*W*^−1^) associated with reaction *j*. To calculate enzyme abundance and turnover number, we followed the procedure outlined in Lewis et al^21^ where we used RNA-Seq gene expression from our 49 samples, and disregarded any steps involving mutation data, as our study does not include such data.

## 2.6. Statistics

We used the Pearson correlation coefficient to analyze the correlation coefficient between continuous variables. For logistic regression analysis, we utilized the liblinear solver in Python with a maximum iteration of 100,000, ensuring convergence. Non-parametric comparisons between groups were performed using the Wilcoxon rank-sum test via the mannwhitneyu function in Python’s scipy.stats module, and the wilcox.test function in R. To adjust for multiple testing, we used the Benjamini-Hochberg method, controlling for the false discovery rate (FDR). For survival analysis, we performed Kaplan–Meier estimation to compare survival outcomes between 2 groups, with the cohort median value used as a cut-off. Cox proportional hazards regression models were fitted using the glmnet package in R, allowing for multivariable survival analysis. For flux enrichment analysis, we used hypergeometric 1-sided test and FDR correction for multiple testing. All statistical analyses were performed using RStudio (version 2023.06.1+524) and Python 3.11.3. Statistical tests were two-sided unless otherwise stated, and p-values below 0.05 were considered statistically significant across all analysis performed.

### 3. Results

## 3.1. Reaction fluxes are significantly different between responders and non-responders

We identified significant differences in metabolic fluxes between responders and non-responders across multiple objective functions (Figure 2A). Three reactions—Succinate Dehydrogenase (SUCD1m), NADH: Guanosine-5-Phosphate Oxidoreductase (r0276), and Ornithine Transaminase Reversible, Mitochondrial (ORNTArm)—were consistently different among all objective functions (Figure 2B). These reactions are involved in the citric acid cycle, purine catabolism, and urea cycle, respectively. Cytosolic NADPH production was also significantly different between responders and non-responders, suggesting a potential role in modulating immune responses.

**Figure 2:**
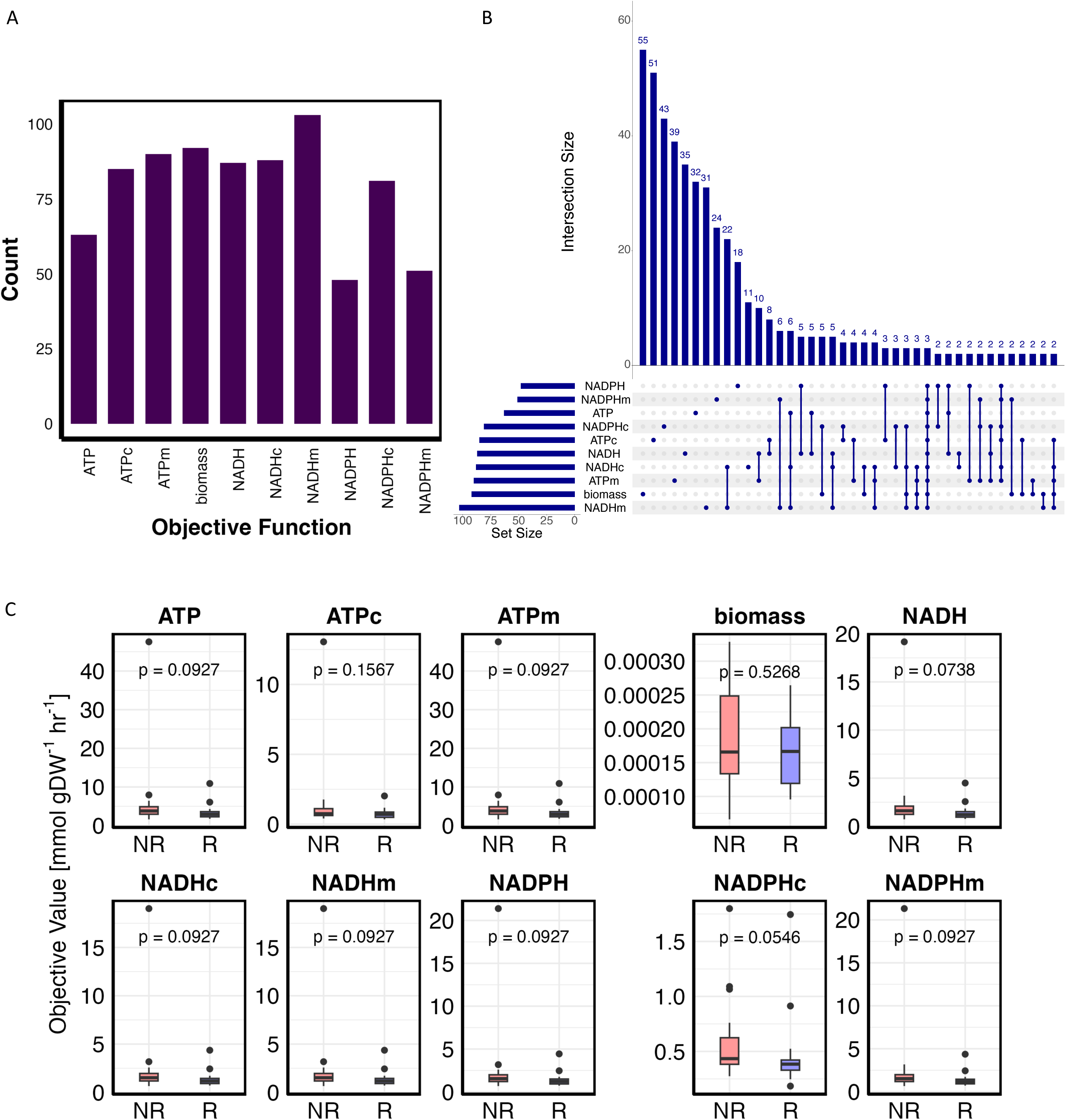
Reaction flux differences across multiple objectives. **(a)** Bar plots representing the count of significantly different reaction fluxes across the ten distinct objective functions. Each bar indicates the number of reactions that exhibit significant differential activity, highlighting the variability in reaction fluxes depending on the specific objective function optimized. **(b)** Upset plot displaying the intersection of mutually significant reactions across the ten different objective functions. The bars quantify the number of reactions that are differentially expressed in each set of objective functions, with the numbers on the bars representing the size of the intersections for each group of metabolites. **(c)** Box plot illustrating the predicted production of key metabolites, including both cytosolic and mitochondrial compartments, for each of the objective functions, including biomass production. The plot includes an indication of the p-value for each comparison, though these p-values have not been adjusted for multiple testing.

The distribution of flux value for these three significant reactions across the different objectives was noted in Supplemental Figure 1A-C. Specifically, responders exhibited lower flux values for both SUCD1m and r0276, with average log2-fold changes of 0.7 and 0.3, respectively. A similar pattern is also observed for ORNTArm, where the average absolute value for responders was lower than in non-responders (log2-fold change of 0.7). In Figure 2C, we compared the FBA-predicted cytosolic and mitochondrial production of each of the ten metabolites, including biomass production. Only the cytosolic NADPH production was close to being significant between responders and non-responders.

We compared the relevant enzyme gene expression for each of these three reactions to show if these differences were also observed in the gene expression data. The result is shown in Supplementary Figure 1D, where we see the four genes (*SDHA*, *SDHB*, *SDHC* and *SDHD*) that support SUCD1m have a higher expression value in non-responders as compared to responders. A similar pattern is observed for r0276, while for ORNTArm which had a higher negative flux for non-responder also showing a higher expression for the associated gene (*OAT)*.

## 3.2. NADH optimizing objective function shows significant reactions in various subsystems

We next focused on the NADH optimizing objective function as it shows a stronger correlation with the immune cell abundance scores derived from our previous publication^1^. The flux enrichment analysis results shown in Figure 3A indicate that fatty acid oxidation subsystem is more enriched for reaction fluxes that are differentially expressed between responders and non-responders. Additionally, the purine catabolism and citric acid cycle subsystems were significantly enriched. We then created a scatter plot in Figure 3B showing the distribution of differentially active reactions between responders and non-responders across various subsystems. To quantify the difference between the mean of the two groups, we also calculated Cohen-d statistic, providing a measure in terms of standard deviation.

**Figure 3:**
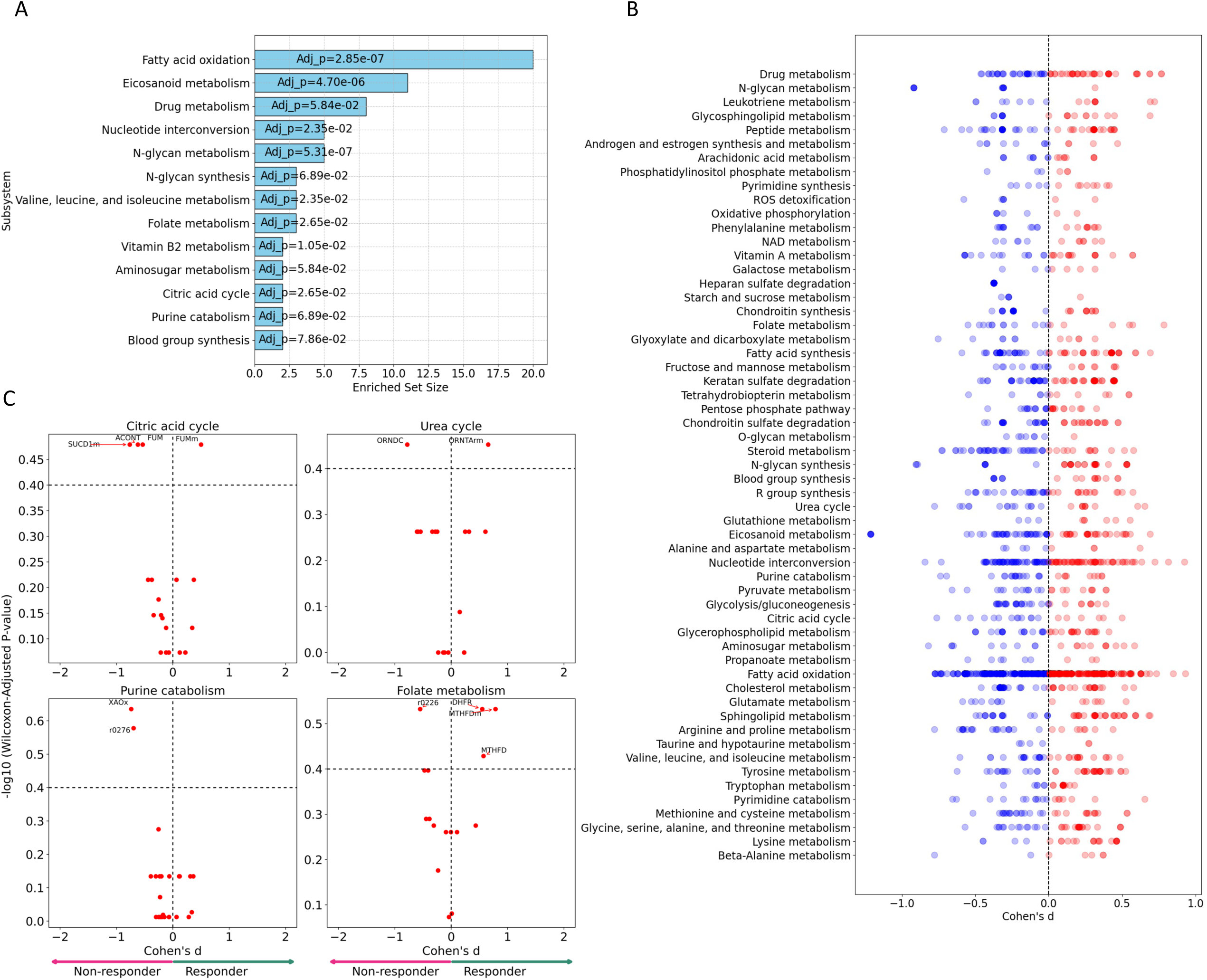
Subsystem analysis using the NADH Objective Optimizing Function. **(a)** Results from Flux Enrichment Analysis showing the enriched subsystems based on differentially expressed reactions using NADH objective optimizing function. The subsystems with significantly enriched reactions are shown, with the adjusted p-values calculated using the False Discovery Rate (FDR) correction method. **(b)** Scatter plots illustrating the differential activity of metabolic reaction fluxes under the NADH objective function. The reactions are grouped by their respective subsystems and color-coded according to the sign of the Cohen’s d statistics, indicating the magnitude and direction of the effect size. Reactions with deep colors signify those with p-values less than 0.05. **(c)** Volcano plot representing differentially expressed reactions in the folate metabolism subsystem, along with the subsystems for the three significant reactions that were differentially expressed across all objectives. The plot highlights both the magnitude of the effect, comparing responders to non-responders and also shows the statistical significance, with p-values adjusted for multiple testing using the Benjamini-Hochberg correction method.

By analyzing the subsystems to which SUCD1m, r0276 and ORNTArm belong, we identified additional reactions that are differentially expressed between responders and non-responders. As shown in Figure 3C, within the urea cycle subsystem, both ORNDC (Ornithine Decarboxylase) and ORNTArm showed significant difference in flux distribution between responders and non-responders. ORNDC had a negative Cohen’s d value, indicating that its average flux is higher in non-responders compared to responders, while ORNTArm showed the opposite trend. In the citric cycle subsystem, we observed that ACONT (Aconitase), FUM (Fumarase) and SUCD1m had higher average flux in non-responders versus responders, whereas FUMm (Fumarase, Mitochondrial) had a higher average flux in responders. In addition to r0276, XAOx (Xanthine Oxidase, Peroxisomal) also had a higher average flux in non-responders in the purine catabolism subsystem.

Our result also shows that in the folate metabolism subsystem, r0226 (5, 6, 7, 8-Tetrahydrofolate: NADP+ Oxidoreductase), DHFR (Dihydrofolate Reductase), MTHFDm (Methylenetetrahydrofolate Dehydrogenase (NADP), Mitochondrial) and MTHFD (Methylenetetrahydrofolate Dehydrogenase (NADP)) displayed significant differences in activity between responders and non-responders.

## 3.3. SUCD1m is highly positively correlated with CD47 expression and TGF-β score

In our previous studies^1,12^, we provided insight into the potential role of *CD47* and TGF-β in mediating immunotherapy resistance. We showed that higher *CD47* levels were associated with lower response to ICI and trended toward lower PFS in the same gynecological cancer cohort. Additionally, elevated *CD47* levels were associated negatively with PDL1 and *CTLA4* expression, as well as cytotoxic T-cells and dendritic cells but positively with TGF-β, BRD4 and CXCR4/CXCL12 expression. Here, we sought to find the correlation between the three reactions flux and *CD47* and TGF-β. Interestingly, we found that SUCD1m is both significantly positively correlated with both *CD47* and TGF-β when optimizing for NADH production (Figure 4A), while ORNTArm had a significant negative correlation with both *CD47* and TGF-β (Supplementary Figure 2A). No significant correlation was observed between r0276 and either *CD47* or TGF-β (Supplementary Figure 2B). The correlation results with other immune cell abundance scores are given in Figure 4B. We observe that, in addition to *CD47* and TGF-β, SUCD1m is also positively correlated with *CD276*, but negatively correlated with other immune cell types. Of interest are the reactions included in folate metabolism subsystem, MTHFD2(m), MTHFD(m) and DHFR. The result shows that MTHFD2m (Methylenetetrahydrofolate Dehydrogenase (NAD), Mitochondrial) is significantly positively correlated with almost all of the immune cell types, while MTHFDm (Methylenetetrahydrofolate Dehydrogenase (NADP), Mitochondrial) showed significant negative correlation with *CD274*, *CTLA4*, plasma cells, dendritic cells and mast cells.

**Figure 4:**
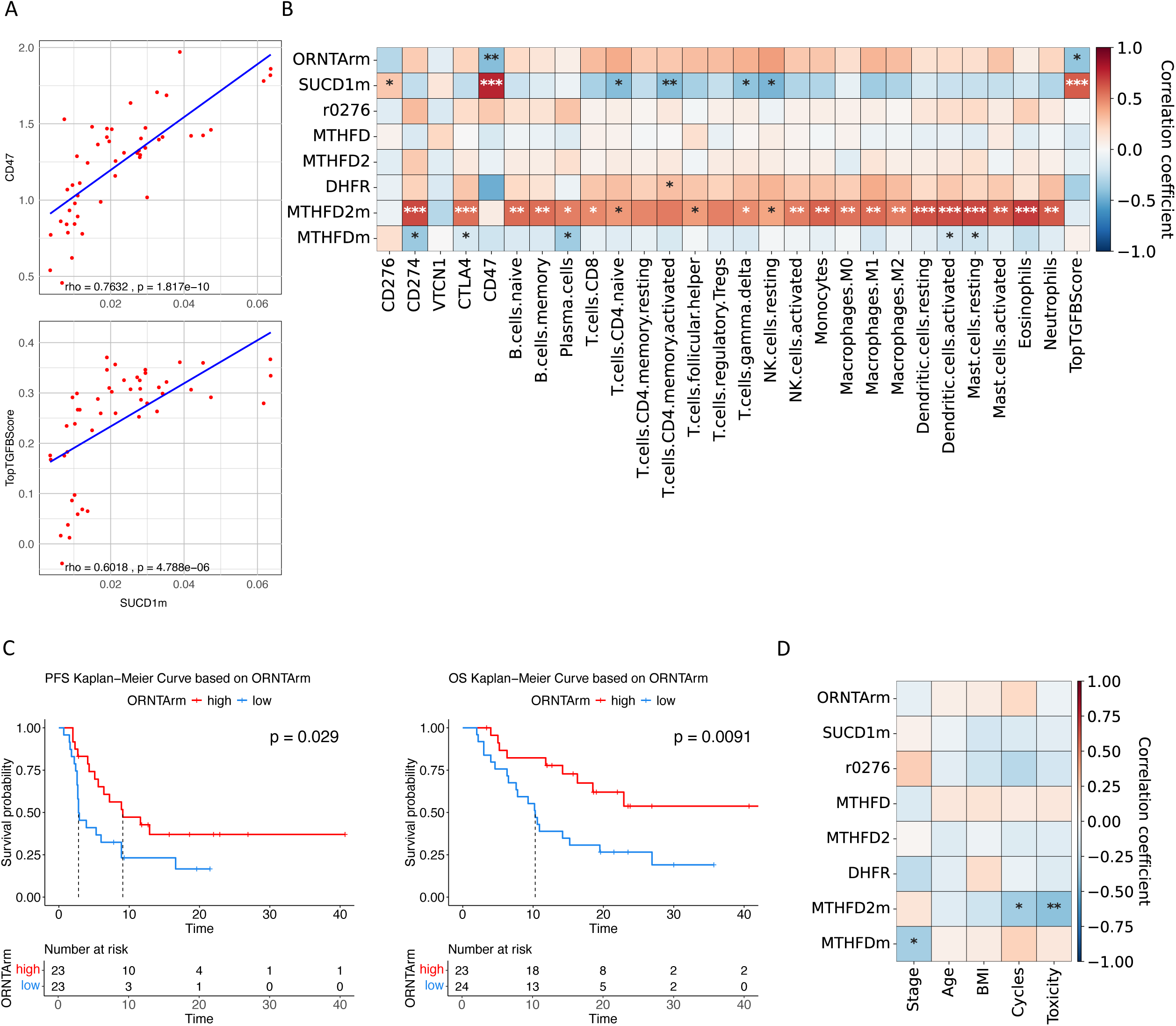
Correlation between highlighted metabolic reaction fluxes and previously reported molecular markers using NADH optimizing objective. **(a)** Scatter plot showing the relationship between of SUCD1m flux and two key molecular markers, CD47 and TGF-β under the NADH optimization objective function. The plot illustrates how SUCD1m correlates with these immunosuppressive molecules, giving an insight into metabolic influence on the immune response in the tumor microenvironment. **(b)** Correlation heatmap showing Pearson’s correlation coefficients between molecular features and selected metabolic reactions, including the three significant reactions identified. This heatmap illustrates the magnitude and direction of correlations between metabolic fluxes and molecular features. **(c)** Kaplan-Meier curves for Progression Free Survival (PFS) and Overall Survival (OS) based on ORNTArm reaction flux under the NADH objective function. The survival plots compare patients with high versus low flux in ORNTArm, where the median value is used to separate between high and low flux. This plot illustrates the impact of metabolic activity on patient survival. **(d)** Pearson’s correlation coefficient heatmaps between clinical features and selected metabolic reaction fluxes under the NADH objective function.

## 3.4. There is significant difference in survival between patients with low-high ORNTArm flux

We analyzed the survival outcomes of patients based on the flux levels of ORNTArm, focusing on their Progression-Free Survival (PFS) and Overall Survival (OS). To categorize patients into low and high flux groups, we used the median flux value as a threshold. Our analysis revealed a significant difference in survival outcomes for both PFS and OS based on ORNTArm flux levels (Figure 4C). Specifically, patients with higher ORNTArm flux demonstrated significantly better survival outcomes for both PFS and OS compared to those with lower flux levels. In contrast, when we examined the flux levels of SUCD1m and r0276, we did not observe any significant differences in survival outcomes for either PFS or OS, as shown in Supplementary Figure 3. Interestingly, these 3 reactions were not correlated or significantly different based on other demographic-clinical features such as stage, age, BMI (Figure 4D). MTHFD2m reaction, on the other hand wass associated with the cycles of ICI and consequent toxicity.

## 3.5. Fluxomic features are important in predicting response to immunotherapy

We developed a logistic regression classifier to assess the predictive value of these multi-omics features in determining responses to immunotherapy. Our model incorporated clinical-demographic (age, cancer type, stage) and molecular-genomic features, including immune marker gene expression (*CD276, CD274, VTCN1, CTLA4* and *CD47*), TGF-β score, 22 immune cell abundance from transcriptomic deconvolution, and the reaction fluxes of SUCD1m, ORNTArm, r0276, MTHFDm and MTHFD2m.To account for missing data, we excluded three samples with missing values for ‘Stage of disease’, thus reducing the sample size from 41 to 38. We split our data into the testing and training sets, with the testing set comprising 5 samples, and the training set 33 samples. Given the small dataset size, we utilized GridSearchCV with StratifiedKFold cross-validation. Our optimal model was an L1-regularization model with a C-value of 0.4329. The advantage of logistic regression with L1-regularization is its ability to identify features that are important in the classification task. As shown in Figure 5A, three metabolic features (MTHFDm, SUCD1m and ORNTArm) emerged as significant in distinguishing between responses to immunotherapy. To validate the reliability of the result, we also present the confusion matrix in Figure 5B, which shows 80% accuracy on the test set.

**Figure 5:**
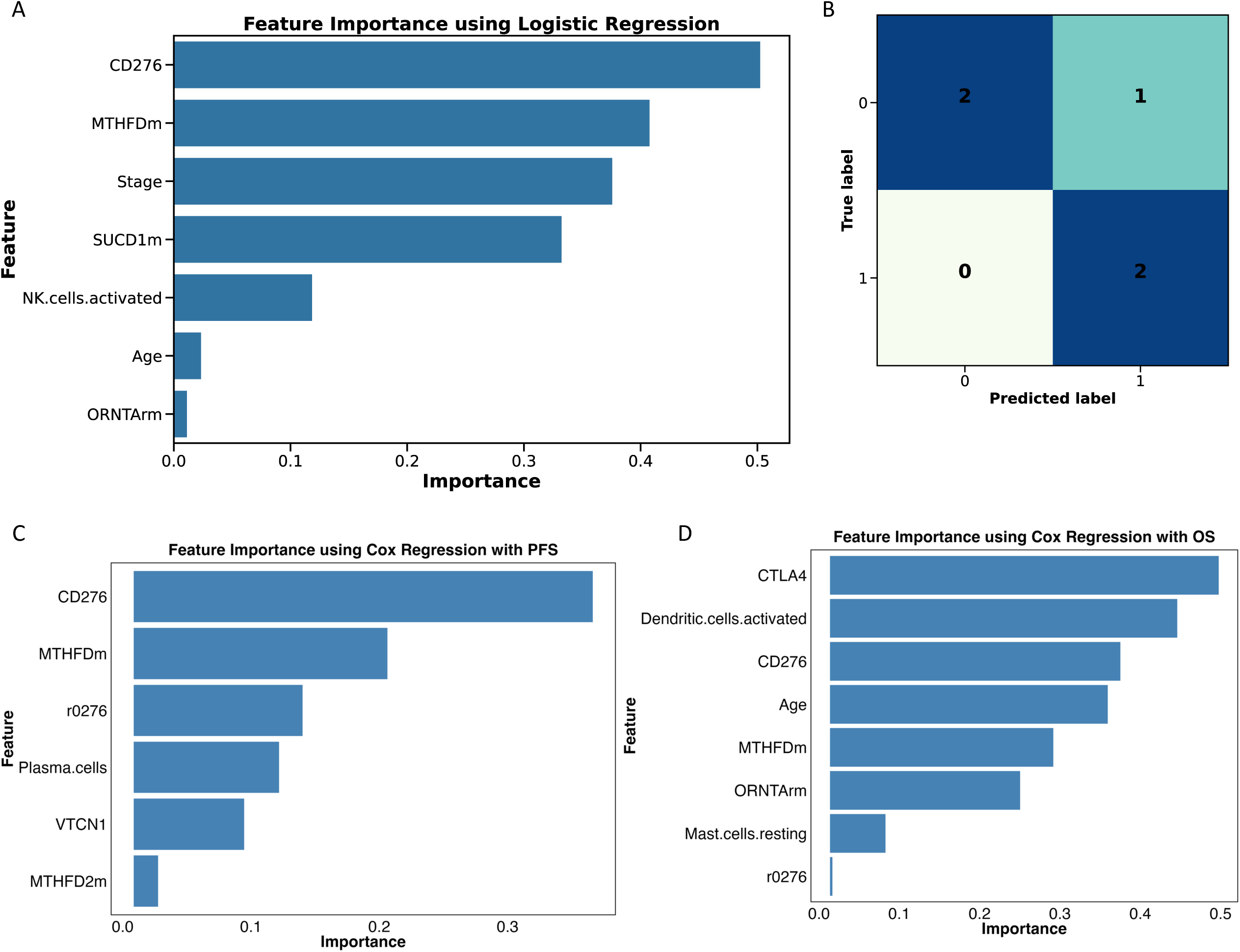
Feature importance using logistic regression and regularized cox model. **(a)** Feature importance using Logistic Regression model for predicting treatment response. This plot shows the features most influential in the model to determine whether a patient will respond to treatment, ranked by their importance. **(b)** Confusion matrix for the 5-sample test set, showing the performance of the Logistic Regression model. The matrix represents the number of true positives, true negatives, false positives and false negatives, which helps to evaluate how best our model is performing. **(c)** Feature importance from regularized cox model using progression and PFS data. The plot shows key features that impact the prediction of disease progression, ranking them from top to bottom. **(d)** Feature importance from regularized cox model using death and OS. Similar to 4C, the plot shows the important factors predicting patient mortality.

Additionally, we developed a regularized cox model to examine the relationship between predictor variables and OS or PFS (Figure 5C-D). This analysis highlighted CD276, MTHFDm, and r0276 as the top predictors for disease progression, while CTLA4, dendritic cells, CD276, Age, and MTHFDm were identified as key predictors for patient mortality.

### 4. Discussion

In this study, we developed personalized genome-scale models to investigate metabolic variations and systems-wide changes in redox metabolism among GYN cancer patients who received immunotherapy. By employing an integrative approach that combines multi-omics data in the model development and machine learning pipeline, we were able to identify key metabolic pathways that differentiate responders from non-responders to immune checkpoint inhibitors. The use of GSMM and FBA allowed us to identify key metabolic pathways that differ between responders and non-responders of ICI, providing insight into potential mechanisms of resistance. The strong correlations between specific metabolic reactions and immune markers, such as CD47 and TGF-β, further emphasize the importance of redox metabolism in modulating immune responses. As a positive control for our analysis, reactions within the folate metabolism subsystem, particularly those involving MTHFD2 have been reported as important markers for distinguishing responders from non-responders across various cancer types. For example, Shang et al^27^ demonstrated that MTHFD2 plays a significant role in cancer immune evasion by upregulating PD-L1 expression, which is critical for tumor growth and immune resistance. They postulated that MTHFD2 may well be a safe therapeutic target for cancer treatment. High MTHFD2 expression has also been linked to an inflamed TME and better responses to ICI in bladder cancer, correlating with increased immune cell infiltration and better survival outcomes^28^. Additionally, in ovarian cancer, recent findings^29^ demonstrate that MTHFD2 overexpression is associated with tumor progression, and its inhibition triggers ferroptosis via ERK signaling, potentially suppressing malignancy. In our work, we showed MTHFD2 to be strongly correlated with stage, cycle and toxicity (Figure 4D), and it was also among the top features for predicting response to ICI (Figure 5A, C, D). Also, our data showed MTHFD2 reaction is highly positively correlated with majority of the immune cell abundance (Figure 4B) and was also among the important features that impact the prediction of disease progression (Figure 5C).

Another important metabolic reaction we found was succinate dehydrogenase (SUCD1m) in the TCA cycle, which oxidizes succinate to fumarate. This reaction flux was upregulated in non-responders as compared to responders (Supplementary Figure 1B). Both succinate and fumarate have been reported as oncometabolites in various cancer types^30–33^. Mutations in succinate dehydrogenase (*SDHx*) genes predispose to hereditary paraganglioma-pheochromocytoma syndrome as well as kidney and gastrointestinal tumors. In our previous study^34^, we reported that germline succinate dehydrogenase gene mutations lead to increased reactive oxidative species (ROS), accumulation of succinate, which stabilized and hyperactivated hypoxia inducible factor (HIF)1α signaling. *SDHx* mutations are also associated with loss of p53, which is regulated by MDM2-independent NADH quinone oxidoreductase 1-mediated protein degradation, likely due to the imbalance of flavin adenine dinucleotide (FAD)/nicotinamide adenine dinucleotide (NADH) caused by SDH alteration. In one of our follow up studies^35^, we identified fumarate as a pertinent metabolite, distinguishing individuals who develop autism spectrum disorder (ASD) from those who develop cancer, with implications for potential biomarker and/or therapeutic value. Similarly, our findings show that the reaction SUCD1m, which converts succinate to fumarate had higher flux in non-responders, suggesting elevated succinate levels and reduced fumarate levels in responders as compared to non-responders. Furthermore, the strong positive correlation of SUCD1m flux with previously reported biomarker of *CD47* and TGF-β (Figure 4A) underscores its potential as a metabolic biomarker for understanding resistance mechanisms to ICI in gynecological cancers.

The integration of transcriptomics data into GSMMs enables generation of context-specific GSMMs and prediction of context-specific metabolic fluxes tailored to specific conditions. Our approach for achieving this differs significantly from traditional approaches which are based on model extraction methods (MEMs) such as iMAT, GIMME, FASTCORE, INIT, and others. The core strategy of these MEMs is to utilize an arbitrary gene expression threshold to remove reactions associated with low gene expression levels. The shortcoming of these approaches is that there is no clear correlation between gene expression level and the expression of proteins that catalyze a reaction, therefore important reactions may be excluded from the reduced model. In our study, rather than removing reactions based on gene expression level, we have integrated both kinetic and thermodynamic data with transcriptomics data into constraints on the maximum metabolic fluxes and the directionalities of individual reactions, reducing the feasible solution space and thereby improving prediction accuracy. This incorporation of kinetic or thermodynamic constraints has been shown to greatly affect metabolic fluxes and directionalities of individual reactions^36^. Additionally, our approach is unique in that we have implemented an objective-customizable genome scale model. The choice of the appropriate objective that cells aim to optimize is still a controversial issue. While most studies assume that cells aim to optimize biomass production, we hypothesize that mammalian cells could have different or multiple objectives depending on environmental conditions, growth phases, and metabolic needs. Instead, rather than basing our prediction on a single metabolic objective, we optimize a set of objective functions, such as maximizing redox cofactor reduction and ATP production. By identifying reaction fluxes that are consistently differentially expressed across these objectives, we provide a more comprehensive and accurate representation of metabolic differences between responders and non-responders. This innovative approach offers valuable insights into the metabolic underpinnings of resistance to ICIs and potential therapeutic targets in gynecologic oncology.

Several limitations should be acknowledged in our study. First, the small sample size may limit the generalizability of our findings, as a larger cohort would provide greater statistical power and robustness. Second, the inclusion of mixed gynecological cancer types introduces heterogeneity, which may impact the interpretation of metabolic patterns across specific cancer subtypes. Future work will focus on expanding the sample size to improve statistical significance and stratify results by specific gynecological cancer subtypes. Additionally, our analysis relies on static flux balance analysis (FBA), which, while informative, does not account for dynamic parameters such as temporal changes in metabolism or cellular adaptation over time. To address this limitation, we are currently incorporating dynamic modeling approaches to better capture the temporal and adaptive nature of metabolic reactions identified in our current analysis.

### 5. Conclusion

We investigated whether there are significant fluxomic biomarkers that distinguish between responders and non-responders to immune checkpoint inhibitors in a cohort of gynecologic cancers, including ovarian, cervical, and endometrial cancers. Using an objective-customizable flux balance analysis to simulate a genome scale metabolic model, we found that three reactions, Succinate Dehydrogenase (SUCD1m) which is found in the citric acid cycle subsystem, NADH: Guanosine-5-Phosphate Oxidoreductase (r0276) involved in purine catabolism, and Ornithine Transaminase Reversible, Mitochondrial (ORNTArm) in the urea cycle were consistently different among all objective functions. Our model also identified certain reactions in the folate cycle subsystem such as MTHFD2 which has been reported to be significant between responders and non-responders. Furthermore, we developed machine learning models to study the relationship between predictor variables and survival time and to investigate whether there are multi-omics biomarkers that are associated with resistance to ICI, which can help to further understand mechanisms driving resistance to ICIs in the field of gynecologic oncology.

In summary, if validated in a larger cohort, these metabolic biomarkers could potentially lead to the development of novel therapies to improve the management of gynecological cancers.

### Data availability

The RNA sequencing data used for this study are deposited in the SRA repository (SRP341153) with the following accession number: PRJNA770873.

### CRediT authorship contribution statement

GI, HM, and YN conceptualized the study. GI and LL carried out the model perdition data analyses and/or performed statistical analyses. GI, HM, and YN interpreted the analyses. GI and YN drafted the manuscript. GI, LH, HM, and YN critically revised the manuscript. YN oversaw the project. All authors read and approved the final manuscript.

---

**Supplemental Table 1:** Clinico-pathological summaries of 49 gynecology patients.

|  |  |
| --- | --- |
| Features | N = 49 |
| **DiseaseSite** |  |
| Ovarian | 14 (29%) |
| Endometrial | 27 (55%) |
| Cervical | 8 (16%) |
| **StageAtDiagnosis** |  |
| I | 13 (29%) |
| II | 2 (4.4%) |
| III | 17 (38%) |
| IV | 13 (29%) |
| NA | 4 |
| **MSI** |  |
| Stable | 16 (33%) |
| High | 17 (35%) |
| NA | 16 (33%) |
| **Age** |  |
| Median (IQR) | 68 (62, 74) |
| NA | 1 |
| **Cycles** |  |
| Median (IQR) | 8 (3, 11) |
| **Toxicity** |  |
| 1 | 23 (48%) |
| 0 | 25 (52%) |
| NA | 1 |
| **Response** |  |
| Progressive disease | 21 (45%) |
| Responded disease | 20 (43%) |
| Stable disease | 6 (13%) |
| NA | 2 |
| **Death** |  |
| 1 | 29 (60%) |
| 0 | 29 (60%) |
| NA | 1 |
| Overall Survival (months) |  |
| Median (IQR) | 13 (6, 21) |
| NA | 1 |
| Progression |  |
| 1 | 33 (72%) |
| 0 | 13 (28%) |
| NA | 3 |
| Progression-Free Survival (months) |  |
| Median (IQR) | 6 (3, 12) |
| NA | 2 |

**Figure legends**

**Supplementary Figure 1:** Distribution of fluxes for the reaction **(a)** r0276, **(b)** ORNTArm and **(c)** SUCD1m across the different objectives.

**Supplementary Figure 2:** Scatter plot showing the relationship between ORNTArm and r0276 fluxes and two key molecular markers, CD47 and TGF-β under the NADH optimization objective function.

**Supplementary Figure 3:** Kaplan-Meier curves for Progression Free Survival (PFS) and Overall Survival (OS) based on SUCD1m and r0276 reaction fluxes under the NADH objective function.

