## Supplementary figures and images for "Metabolic Signatures of Immune Checkpoint Inhibitor Response in Gynecologic Cancers: Insights from Flux Balance Analysis"

### Supplemental Figures

#### Supplementary Figure 1

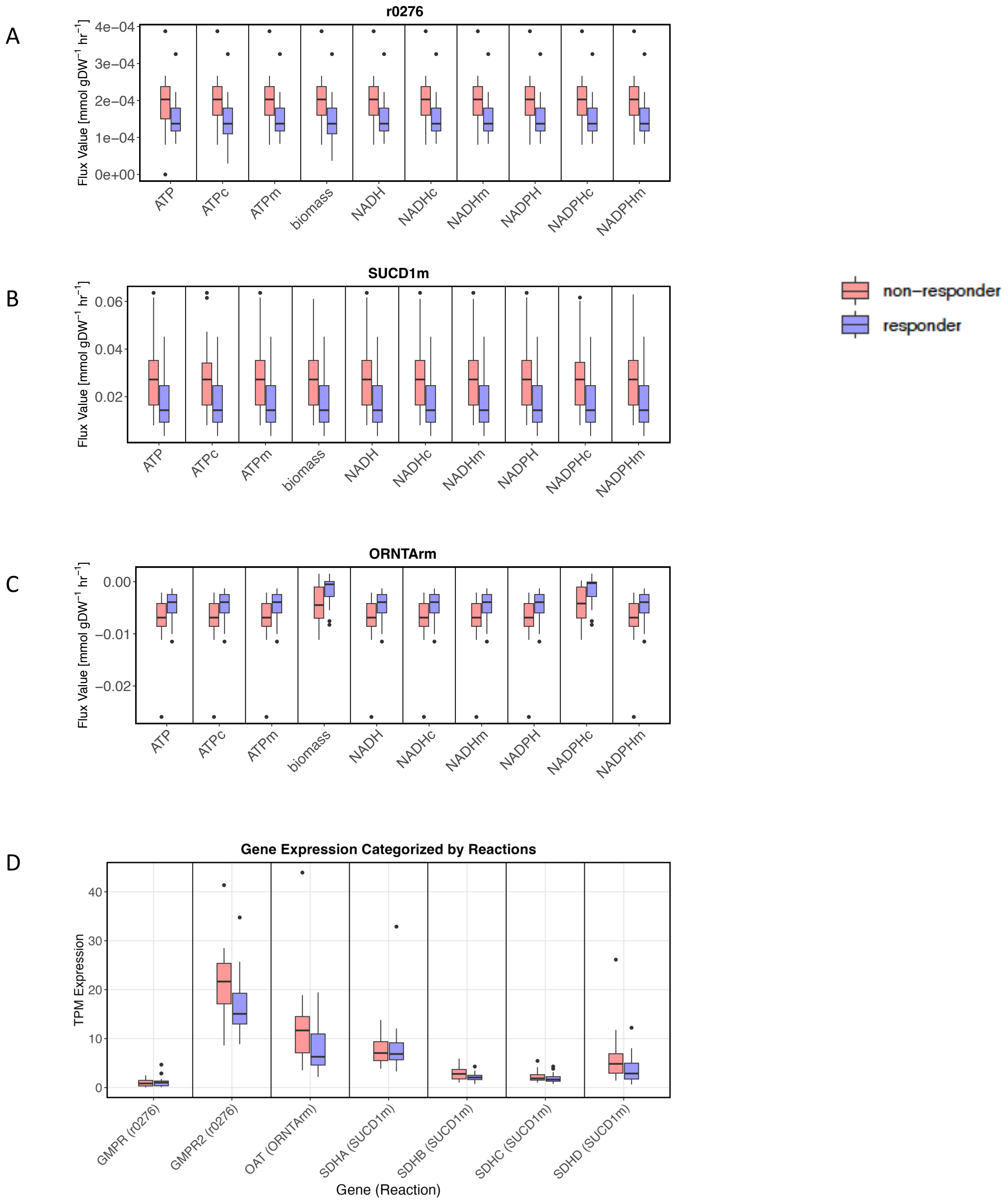

#### Supplementary Figure 2

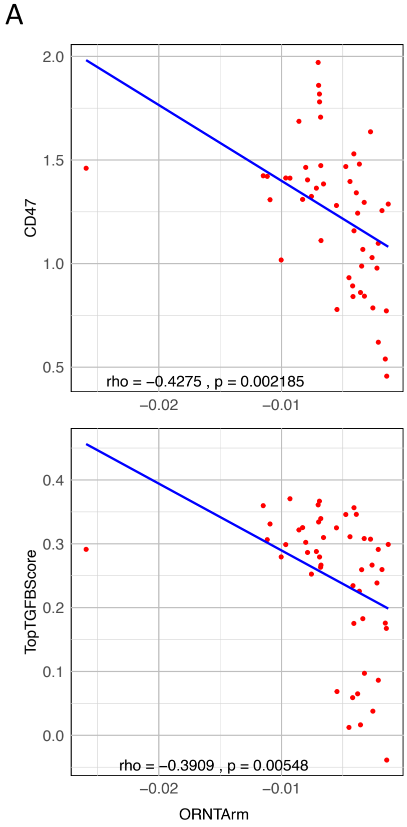


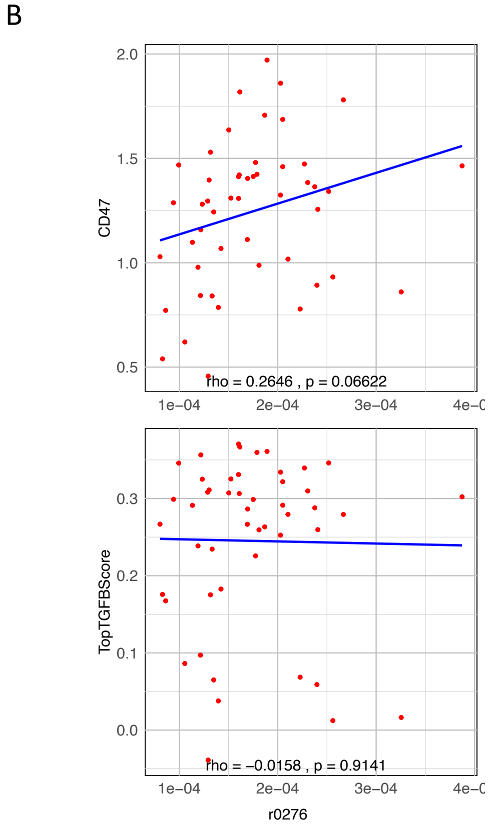

#### Supplementary Figure 3

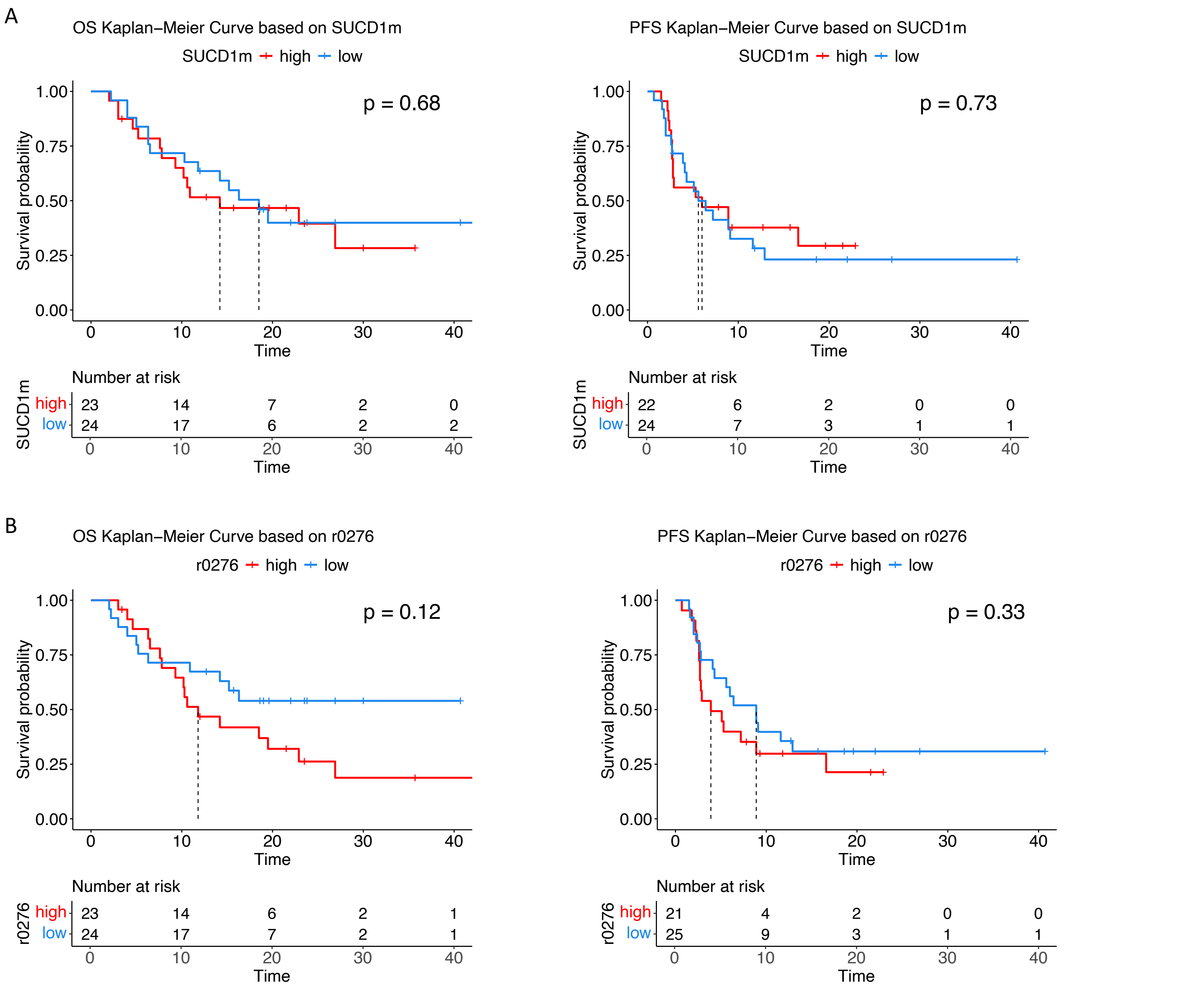
